# Transcriptomic profiling of embryo-derived cell lines from the Chagas disease insect vector *Rhodnius prolixus*

**DOI:** 10.64898/2026.05.08.723764

**Authors:** Laura de Andrade Tavares, Anna Carolina Silva Garcia, Lesley Bell-Sakyi, Tarcísio Fontenele de Brito, Attilio Pane

## Abstract

*Rhodnius prolixus* is a primary insect vector of *Trypanosoma cruzi*, the causative agent of Chagas disease, a neglected parasitosis endemic in Latin American countries. It has been estimated that Chagas disease affects 7-8 million people worldwide and is responsible for approximately 1000 deaths per year. Genetic and molecular studies in this species remain challenging due to its life cycle and feeding habits, thus hindering the development of new strategies to control their populations and reduce the diffusion of Chagas disease. Recently, two stable cell lines - RPE/LULS53 and RPE/LULS57 - were derived from *Rhodnius* embryos, which represent promising new tools to investigate the genetics of this insect vector. Here, we describe their gene expression landscapes through transcriptomic approaches. We show that 8,968 expressed genes are shared between the two cell lines, whereas 391 and 1,088 genes are uniquely expressed in RPE/LULS53 and RPE/LULS57, respectively. Although key components of primary developmental, immune and redox signaling pathways are expressed in both cell lines, some genes such as *Frizzled-10-a-like* and *catalase* show marked differences in expression. Our results strongly suggest that RPE/LULS53 and RPE/LULS57 likely represent two different cell phenotypes. Consistent with this, gene ontology analysis reveals that RPE/LULS53 is enriched for animal organ morphogenesis and stress response, while RPE/LULS57 for DNA-directed RNA polymerase activity, among others. Despite these differences, both cell lines express comparable levels of transcripts from resident transposable elements, including the highly abundant *Mariner* and *LINE/I* elements, as well as horizontally transferred transposons. Our findings shed light on the nature of the RPE/LULS53 and RPE/LULS57 embryo-derived cell lines and provide valuable transcriptomic resources for future genetic and functional studies in *Rhodnius* and other triatomine insect vectors.

**Author summary:** *Rhodnius prolixus* is a blood-feeding insect and a major vector of Chagas disease, a parasitosis endemic in Latin America and affecting millions of people worldwide. In the absence of effective drugs and vaccines, the control of the insect population represents a promising strategy to reduce the diffusion of the disease. Yet, genetic and functional studies in *Rhodnius* are extremely challenging due to its feeding habit and life cycle. To overcome these limitations, researchers have previously developed two stable cell lines derived from *Rhodnius* embryos. In this study, we provide the first characterization of the genes expressed in these cell lines. We found that, while the two cell lines share many expressed genes, each of them also has distinct gene expression patterns pointing to two different cell types with specialized functions. These differences likely affect the way they respond to stress and regulate biological processes. Our findings provide an important resource for researchers studying *Rhodnius prolixus* and other insect vectors, helping advance our understanding of the genetic and molecular mechanisms that control the insect development and mediate the interactions between insect vectors and the parasites they transmit

## Introduction

*Rhodnius prolixus* is a hematophagous hemipteran insect belonging to the family Reduviidae and subfamily Triatominae and is one of the main vectors of the kinetoplastid protozoan parasite *Trypanosoma cruzi*. This parasite is the etiological agent of Chagas disease, a debilitating and potentially fatal illness that predominantly affects socioeconomically vulnerable populations in continental Latin America^1^. First described by Carlos Chagas in 1909, Chagas disease remains one of the most neglected tropical diseases worldwide^2^. According to the World Health Organization (WHO), this parasitosis affects more than 7 million people globally and causes approximately 10,000 deaths annually, posing a substantial public health threat. In response to the persistent impact of neglected tropical diseases, the WHO launched a roadmap for 2021-2030 aiming to strengthen and coordinate global efforts towards their control and elimination^3^.

Beyond its epidemiological relevance, *Rhodnius* has emerged as an important model system for studies of insect physiology, development and vector biology^4–6^. As a hemimetabolous insect, it undergoes incomplete metamorphosis, developing through a series of nymphal stages that progressively resemble the adult form. This contrasts with holometabolous insects such as *Drosophila melanogaster*, which develop through distinct larval and pupal stages before reaching adulthood. Hemimetabolous insects comprise around 10-20% of all insect species, representing phylogenetically older and structurally diverse groups that are essential for understanding the evolutionary origins of insect development and physiology. Despite their medical importance, triatomine insects remain comparatively understudied at the molecular and cellular levels relative to classical insect models.

Among the physiological processes that make *Rhodnius* a particularly informative model, blood feeding is central to its development because each nymphal molt requires a blood meal. As a result, every nymphal stage has the potential to transmit Chagas disease once infected^7^. In adults, blood feeding is critical to activate vitellogenesis and egg production in females. In this insect species, oogenesis occurs in telotrophic ovarioles where trophocytes (i.e. nurse cells) supply macromolecules to the growing ovarian follicle through specialized trophic cords. Following blood feeding, increases in juvenile hormone and ecdysteroids regulate vitellogenin synthesis, oocyte growth, follicular differentiation, and chorion formation^8^. After fertilization, embryogenesis proceeds through rapid syncytial divisions that form a syncytial blastoderm, followed by cellularization, formation of the ventral germ anlage, segmentation, germ band retraction, dorsal closure, organogenesis and cuticle secretion^4,9^. Postembryonic development spans five blood dependent nymphal instars characterized by gradual growth, wing development, and internal maturation without metamorphosis.

Although *R. prolixus* has emerged as a valuable model system, it remains experimentally limited compared with the well-established genetic model system *D. melanogaster*. Stable transgenesis and mutant lines are not currently available and genome editing approaches, such as CRISPR/Cas9, have only recently begun to be explored^10^. In addition, the relatively long life cycle of this insect, which spans approximately six months from embryo to adult, makes multigenerational genetic experiments technically challenging. These limitations highlight the need for complementary experimental systems that enable functional and molecular investigations under more accessible laboratory conditions. A growing number of genomic, transcriptomic and metatranscriptomic studies are starting to shed light on the genetic programs that control *Rhodnius* development, guarantee oogenesis and adult fertility and regulate its interaction with microbes^11–20^.

Recently, the cell lines RPE/LULS53 and RPE/LULS57 were generated from *R. prolixus* embryos providing the first stable cell lines from a hemipteran blood-sucking insect^21^. Insect cell lines have been extensively used to gain a deeper understanding of evolutionarily conserved regulatory mechanisms and gene function. For instance, the *D. melanogaster* cell line S2 has been widely used to dissect conserved cellular pathways, including those governing cellular metabolism^22^. The S2 cells have enabled major advances through the use of RNAi and contributed to identifying fundamental aspects of the mitotic machinery conserved in higher eukaryotes^23^. Among other holometabolous insects, cell lines derived from the silkworm *Bombyx mori* have provided insights into immune functions and metabolism, expanding our understanding of cellular mechanisms and their applications in biotechnology^24^, while the cell lines Aag2 from *Aedes aegypti* and the RNAi-defective C6/36 from *Aedes albopictus* have greatly contributed to dissecting the biology of arboviruses.

In this scenario, the *R. prolixus* embryo-derived cell lines represent valuable tools for investigating gene function, regulatory networks, signaling pathways and genome stability in a hemimetabolous context. These lines will also help dissect triatomine-virus interactions, a field that is still in its infancy. Indeed, only recently, findings from our research group together with previous studies revealed that triatomines harbor multiple RNA viruses that may influence vector immunity, gut physiology, and the ternary interaction among triatomines, viruses and *T. cruzi*, with potential consequences for parasite transmission and vector competence^12,25,26^. The *Rhodnius* embryo-derived cell lines were subjected to Nanopore SISPA RNA-seq to assess the presence of viral infections, revealing that both cell lines are free of *Triatoma virus* as well as a range of viruses that we have recently described^21^. However, the gene expression landscapes of these cell lines are still unknown. In this study, we performed transcriptomic analyses of the RPE/LULS53 and RPE/LULS57 cell lines to fill this gap by characterizing their gene expression profiles and to provide a useful resource for future genetic, genomic and evolutionary studies in triatomines.

## Methods

### RNA extraction, library construction and sequencing

*Rhodnius prolixus* embryo-derived cell lines RPE/LULS53 and RPE/LULS57 were obtained from the Tick Cell Biobank (University of Liverpool) via the South American Tick Cell Biobank Outpost (IOC/Fiocruz). Both cell lines were derived from insects originally collected in Venezuela^21^. Cells were maintained at 28 °C and cultured in 25 cm² TPP culture flasks containing 5 mL of L-15 (Leibovitz) medium supplemented with 20% fetal bovine serum (FBS), 100 U/mL penicillin and 100 µg/ml streptomycin (Thermo Fisher Scientific).

Prior to RNA extraction, the culture medium was discarded and the cells were washed three times with 1× phosphate-buffered saline (PBS) filtered using a 0.2 µm syringe filter. Total RNA was extracted directly from the culture flasks with TRIzol reagent (Invitrogen) according to the manufacturer’s protocol. RNA quality was assessed on a Qubit 4 fluorometer and submitted for library preparation and sequencing at Life Sciences Core Facility (LaCTAD). One replicate was processed for each cell line, yielding approximately 26 million paired-end reads.

### Bioinformatic analysis

Pre-processing of RNA-seq data was performed using snakePipes v2.7.3^48^. Additional parameters (--trim --fastqc --mode alignment-free, alignment, deepTools_qc) were used to enable read trimming, quality control, transcript quantification and alignment. Low-quality reads and adapter sequences were removed with Cutadapt v4.1^49^. Reads were subsequently aligned to the *R. prolixus* genome assembly RproC3.3 (GCA_000181055.3) with STAR v2.7^50^.

Mapped reads were quantified with TEtranscripts v2.2.3^51^, which estimates transcript abundances for both gene and TE. HTT sequences used in this study were previously published^37^. Ribosomal RNAs were filtered using the RproC3.3 annotation as a reference. Raw counts were normalized to TPM and genes with TPM ≥ 1 were retained for downstream analyses.

Gene descriptions were initially extracted from the RproC3.3 gene annotation present in VectorBase. Sequences lacking description were queried against the complete non-redundant protein database (NR) from NCBI (retrieved on August 11, 2025) using DIAMOND blastx v2.1.9.163^52^, retaining hits with e-value ≤ 0.01 and bitscore ≥ 50. Remaining uncharacterized genes were further processed through g:Orth using g:Profiler v0.2.3^53^, with *D. melanogaster* specified as the target organism. Genes without identified orthology or annotation evidence remained as “not assigned” (NA).

GO enrichment analysis was performed using g:GOSt from g:Profiler using the “highlight” parameter, in order to narrow down the most significant findings. Prior to enrichment, genes were filtered with log_10_(TPM + 1) > 1. For expression profiling of developmental, stress and immunity-related pathways, a curated set of genes was selected and is detailed in S9 Table. Similarly, the genes included in the redox homeostasis analysis are listed in S10 Table.

### Molecular cloning and immunofluorescence assays

Genomic DNA was extracted with a genomic DNA extraction kit (New England Biolabs) from 30 first-instar *R. prolixus* nymphs. To isolate the promoter region of the *Rp*-*β-Tub* gene (RPRC004762), a ≈500 bp fragment spanning the putative promoter was amplified with specific oligonucleotides bearing a KpnI restriction site at their 5’-ends.

The resulting amplicon was first cloned into the TOPO-ta plasmid (Invitrogen), generating the TOPO-*Rp*-*β-Tub* construct. Approximately 5 µg of plasmid obtained from a minprep was digested with the KpnI enzyme (New England Biolabs) to release the ≈500bp *Rp*-*β-Tub* fragment. Following gel purification, this fragment was cloned upstream of the eGPF coding sequence in a pBluescript plasmid backbone, producing the pBlue-*Rp*-*β-Tub*-eGFP recombinant plasmid. The appropriate orientation of the putative *Rp*-*β-Tub* promoter relative to the downstream eGFP sequence was assessed by subjecting the pBlue-*Rp*-*β-Tub*-eGFP plasmid to PCR assays using the *Rp*-*β-Tub* forward oligonucleotide in combination with a eGFP reverse oligonucleotide. Only clones with the appropriate orientation were selected for subsequent analysis. The oligonucleotides used for PCR assays and cloning are listed below:

*Rp*-*β-Tub_*KpnI_fw: 5’-TTTGGTACCAGGAGAATAACCTGTACT-3’

*Rp*-*β-Tub_*KpnI_rev: 5’-TTTGGTACCATAGCAAATTACAAGTTCTCC-3’

eGFP_rev: 5’-CAGAAGGACCATGTGGTCGCG-3’

For cell transfection assays, 1 µg of the pBlue-*Rp*-*β-Tub*-eGFP plasmid was transfected into ≈1 million of RPE/LULS57 cells using CellFectin reagent (Gibco) for 1h. Three days post-transfection, the cells were fixed in 4% formaldehyde for 10 min at RT, washed with 1X PBS and subjected to immunofluorescence assays as described previously^11^. GFP was detected using a rabbit polyclonal anti-GFP antibody (Novus Biologicals) and DNA was stained with DAPI (Thermo Fisher Scientific). Image acquisition using a Leica SPE confocal microscope was performed at the Unidade de Microscopia ICB UFRJ.

## Results

### Global gene expression profiles of RPE/LULS53 and RPE/LULS57 cell lines

In order to investigate the gene expression profiles of the RPE/LULS53 and RPE/LULS57 cell lines, we produced and sequenced RNA-seq libraries from total RNA. For each library, we obtained 25,832,421 and 25,964,834 reads, respectively, which were processed through quality filtering, adapter trimming and alignment to the reference *R. prolixus* genome RproC3.3. Approximately 85.21% of the raw reads for RPE/LULS53 and 92.21% for RPE/LULS57 were mapped to the genome (S1 Table).

To characterize the transcriptional profile of both cell lines, we examined the 20 most expressed genes in each dataset. The most abundant transcripts in RPE/LULS53 (Fig. 1a) included genes encoding cytochrome c oxidase subunits, chitin-binding proteins, elongation factor 1α, putative poly(A)-binding proteins and 14-3-3-like proteins. Genes encoding ribosomal components and tubulin were also among the highly expressed transcripts, which is consistent with the abundance of rRNAs in eukaryotic cells and the constitutive expression of tubulin genes required for cytoskeletal organization^27,28^. Six of the 20 top expressed genes in this cell line lack known orthologs or annotated functions.

**Fig 1.**
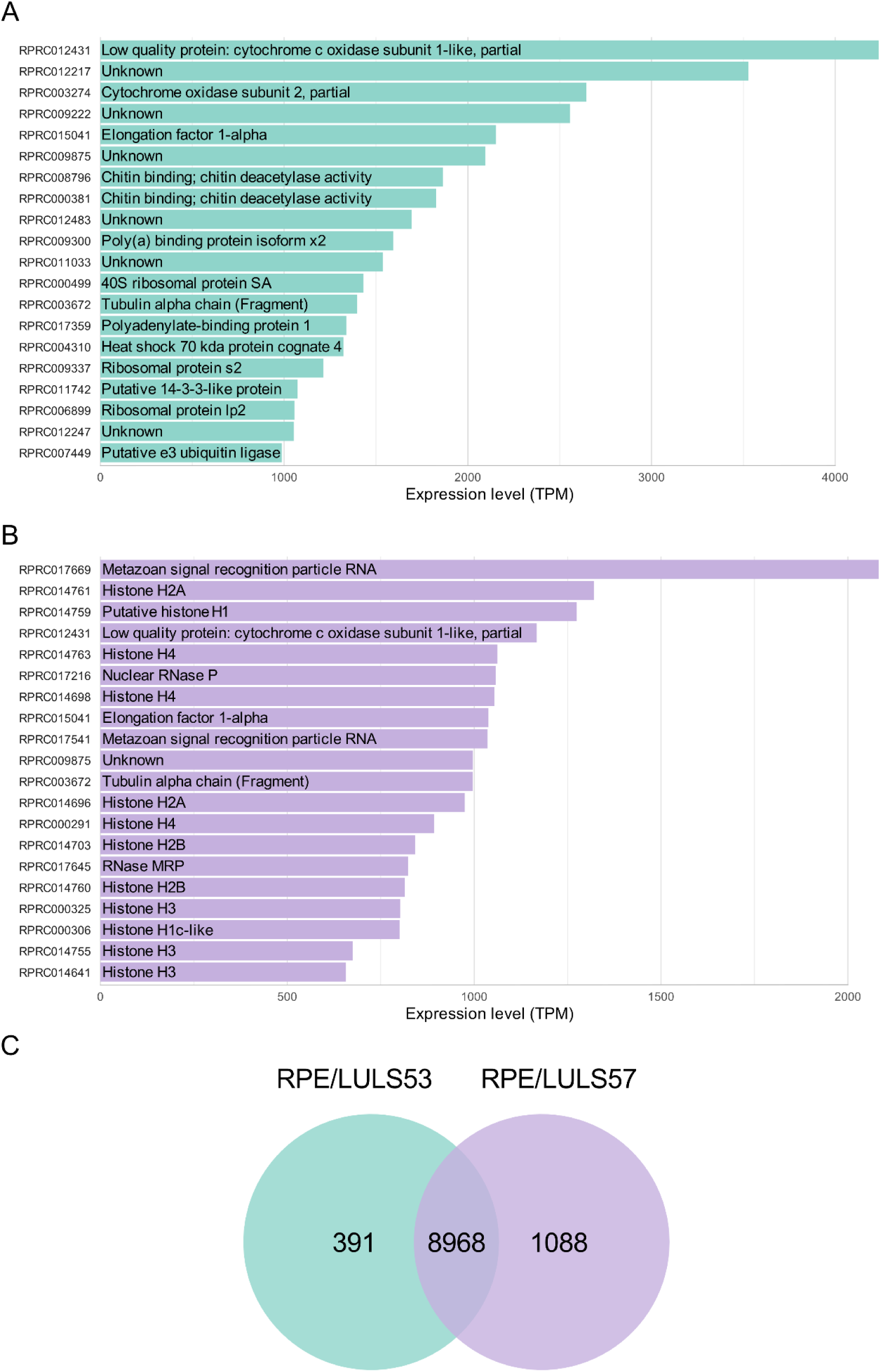
Transcript expression profiles of the RPE/LULS53 and RPE/LULS57 cell lines. (A-B) Bar plots showing the 20 most highly expressed transcripts in RPE/LULS53 (A, green) and RPE/LULS57 (B, lilac), based on RNA-seq data. The x-axis represents the transcript abundance in TPM and the y-axis shows transcript IDs, with the corresponding functional annotation indicated within each bar. (C) Venn diagram illustrating the number of expressed transcripts detected in each cell line applying a TPM > 1 cutoff, highlighting shared and cell line-specific transcripts.

In RPE/LULS57, the most abundant transcript corresponded to the Metazoan signal recognition particle RNA (Fig. 1b), a component of the ribonucleoprotein complex that directs newly synthesized proteins to the endoplasmic reticulum^29,30^. Additionally, 12 of the 20 most highly expressed genes in this cell line encoded histones. Similar to RPE/LULS53, RPE/LULS57 expressed high levels of cytochrome c oxidase subunits, elongation factor 1α and tubulin alpha chain. A complete list of expressed genes (TPM > 1) for each cell line is provided in S2 and S3 Tables.

Gene expression landscapes were compared to highlight similarities and differences between the two cell lines (Fig. 1c). Interestingly, 8,968 genes are shared between the two cell lines, whereas 391 and 1,088 are specific to RPE/LULS53 and RPE/LULS57, respectively.

To further investigate these lineage-specific transcripts, we analyzed the genes uniquely expressed in each dataset. In RPE/LULS53, the 20 top exclusive transcripts were largely composed of genes lacking functional annotation, with no clear orthologs in other species (S1a Fig.). In contrast, the genes expressed only in RPE/LULS57 included several small non-coding RNAs, particularly spliceosomal RNAs of the U1 and U2 classes and small nucleolar RNA U3, as well as multiple genes encoding histone proteins (S1b Fig.). Both spliceosomal RNAs and histone genes belong to multicopy gene families, suggesting that the observed differences between the two cell lines may reflect differential expression of distinct gene copies, a phenomenon that has been shown in different cellular contexts^31^. However, the protocols used for library preparation are not specific for small RNAs and therefore the differential expression of this class of RNAs should be interpreted with caution. Complete lists of genes exclusively expressed in each cell line (TPM > 1) are provided in S4 and S5 Tables.

Together, these results reveal that, despite both cell lines having been derived from *Rhodnius* embryos using similar methodologies^21^, their transcriptional profiles differ at least in part.

### RPE/LULS53 and RPE/LULS57 embryo-derived cell lines are functionally divergent

To assess the functional implications of the differences in their gene expression landscapes, we performed a comparative Gene Ontology enrichment analysis (Fig. 2). While both RPE/LULS53 and RPE/LULS57 revealed enrichment for core gene expression-related categories - including structural constituents of the ribosome, translation factor activity, RNA binding and DNA replication - each cell line displayed additional distinct functional signatures. Interestingly, the highlighted enriched terms were restricted to the Biological Process and Molecular Function categories, with no significant enrichment detected for Cellular Component terms, suggesting that the dominant features of the transcriptome are defined by functional and regulatory activities rather than by compartment-specific features.

**Fig 2.**
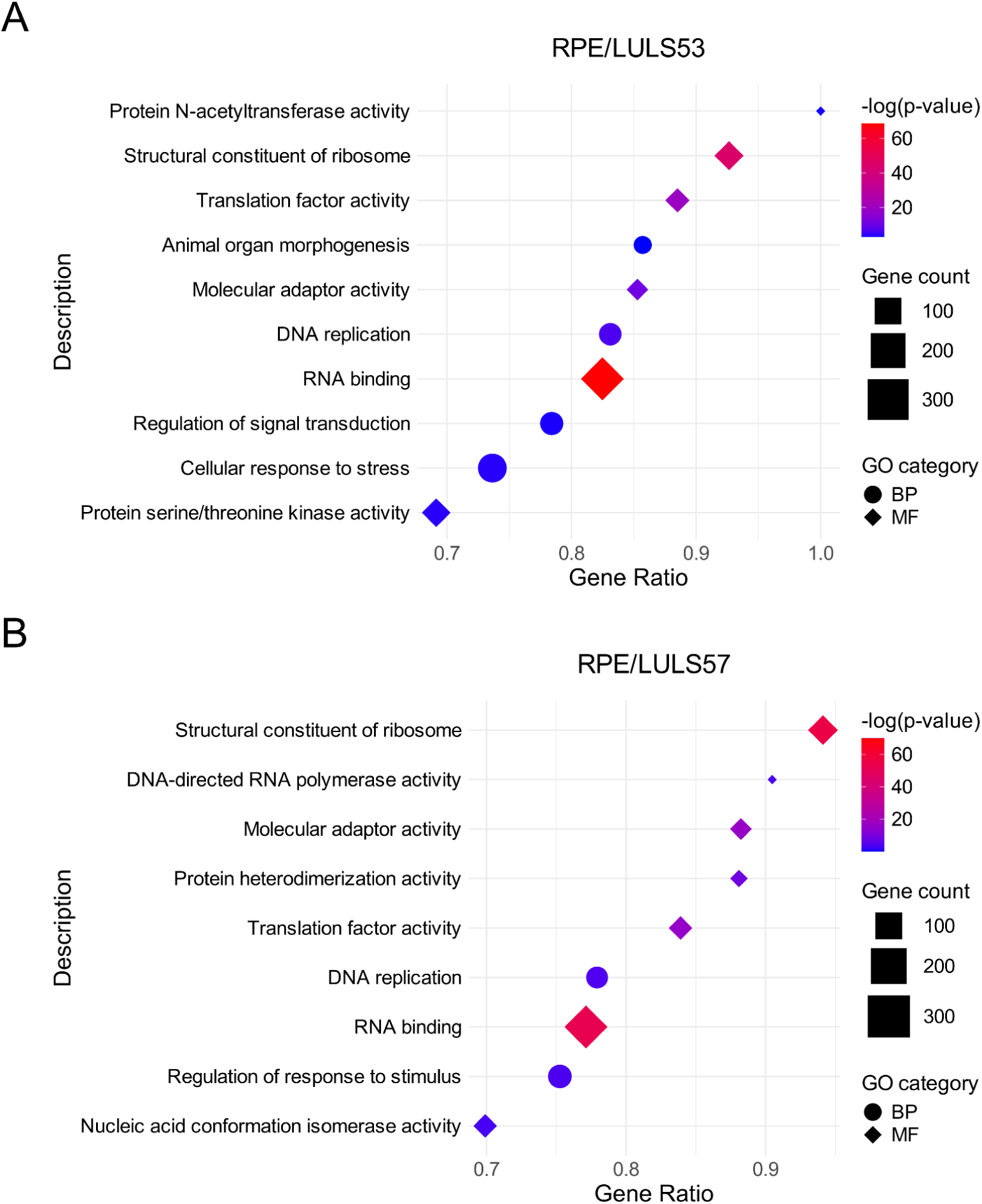
Functional analysis of expressed genes in the RPE/LULS53 and RPE/LULS57 cell lines. Dot plots showing significantly enriched GO terms for (A) RPE/LULS53 and (B) RPE/LULS57. Each point represents an enriched GO term, with the x-axis indicating the gene ratio (proportion of genes in the input list associated with the term) and the y-axis displaying the corresponding functional description. Point size reflects the number of genes associated with each term, while color indicates statistical significance in −log(p-value). Distinct shapes represent the GO categories Biological Process (BP, circles) and Molecular Function (MF, diamonds).

RPE/LULS53 exhibited enrichment for a broad range of biological processes, including regulation of signal transduction, cellular response to stress, animal organ morphogenesis and protein serine/threonine kinase activity (Fig. 2a). These categories point to increased representation of signaling, regulatory and developmental pathways. Meanwhile, enriched terms in RPE/LULS57 were largely restricted to processes associated with transcription, translation and nucleic acid metabolism, such as DNA-directed RNA polymerase activity and nucleic acid conformation isomerase activity (Fig. 2b), a pattern that is associated with a transcriptionally and translationally active cellular state.

Notably, a separate GO enrichment analysis targeting cell line-specific genes revealed that only RPE/LULS57 could be categorized into distinct, biologically relevant functions (S2 Fig.). The enriched terms involved molecular functions, such as structural constituent of chromatin and protein heterodimerization activity, but also cellular component terms, including nucleosome and protein-DNA complex. Conversely, the genes exclusive to RPE/LULS53 failed to produce a robust functional profile, producing only a single, minor category with minimal gene representation.

These enrichment patterns highlight functional differences between the two cell lines, prompting further investigation into specific signaling pathways and regulatory mechanisms. To this end, we examined the expression levels of core components of major developmental signaling pathways, including Wnt, Notch, Bone Morphogenetic Protein (BMP), Hedgehog, Hippo and Planar Cell Polarity (PCP) (Fig. 3a). We further assessed immune-related and stress-responsive pathways, including Toll, Immune deficiency (IMD), Janus kinase/signal transducer and activator of transcription (Jak/Stat) and Extracellular signal-regulated kinase (ERK) (Fig. 3b).

**Fig 3.**
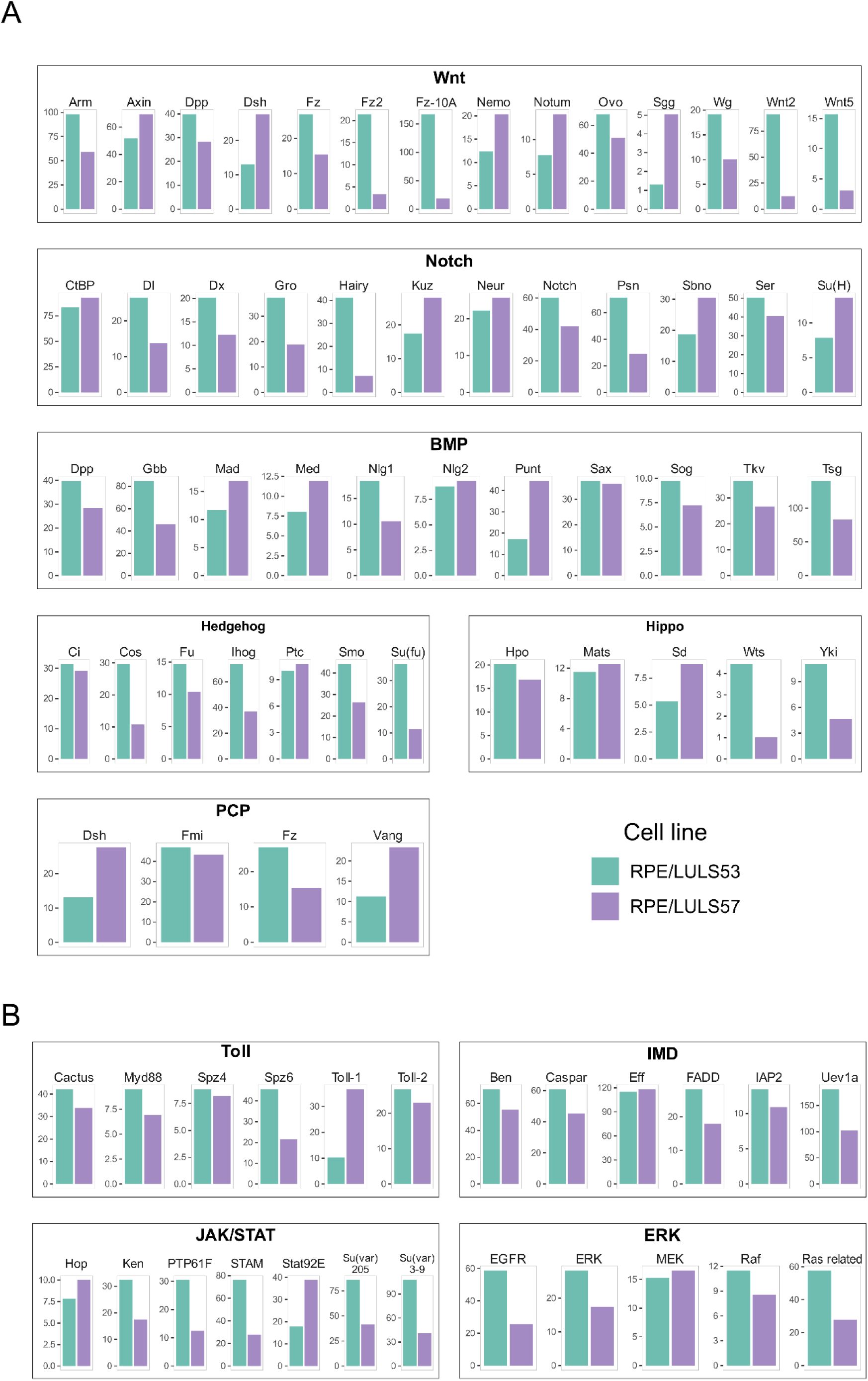
Expression profiles of genes associated with developmental, stress and immune signaling pathways in RPE/LULS53 and RPE/LULS57 cell lines. (A) Expression levels of genes involved in the major developmental signaling pathways Wnt, Notch, BMP, Hedgehog, Hippo and PCP. (B) Expression levels of genes associated with the stress and immune-related pathways Toll, IMD, JAK/STAT and ERK signaling. Gene expression values were derived from RNA-seq data and normalized by TPM. Bars represent transcript abundance displayed for each gene on an independent TPM scale (y-axis).

The comparison indicates that both cell lines retain the core components of several conserved developmental and immune signaling pathways. Nevertheless, quantitative differences in expression levels were observed for several pathway components, although these were exploratory due to the lack of biological replicates. Most genes involved in the Wnt pathway displayed higher expression in RPE/LULS53; for example, the expression of *Wnt2* was over 7 times higher in RPE/LULS53 than in RPE/LULS57 (91.21 vs 12.48 TPM), while *Frizzled-10-a-like* (*Fz-10A*) displayed a 9-fold increase (166.90 vs 18.41 TPM). Moreover, most components of the Hedgehog and IMD pathways also appear to be expressed at higher levels in RPE/LULS53. Conversely, some elements of the Jak/Stat and BMP pathways displayed higher expression in RPE/LULS57, including *STAT92E* (≈2.1 times higher) and the BMP receptor *Punt* (≈2.5 times higher). Overall, these observations indicate that, although the two embryo-derived cell lines preserve a common repertoire of signaling pathways, the relative expression levels of their components differ. The complete dataset of developmental, stress and immune signaling pathways expression levels is available in S6 Table.

Additionally, we investigated the expression of genes involved in heme metabolism and redox homeostasis. This analysis is of particular interest given the hematophagous nature of *Rhodnius*, which ingests large amounts of blood, resulting in a substantial influx of iron and heme. These molecules can promote the generation of reactive oxygen species (ROS), imposing oxidative challenges at the cellular level. In this context, hematophagous insects have evolved mechanisms to prevent iron and heme overload and mitigate oxidative damage^32^. To explore how these processes are regulated, we defined an integrated redox homeostasis network comprising genes involved in ROS metabolism, antioxidant defense, and heme and iron handling. We then analyzed the expression of these components in the RPE/LULS53 and RPE/LULS57 cell lines.

The comparison of redox-related genes revealed additional transcriptional differences between the two cell lines (Fig. 4). Most components of this mechanism displayed higher expression levels in RPE/LULS53 compared to RPE/LULS57. Genes associated in ROS production were consistently more expressed in RPE/LULS53, with *DUOX* (52.52 vs. 20.33 TPM) and *DUOXA1* (26.18 vs. 6.99 TPM) showing some of the most pronounced differences in this category. A similar pattern was observed for antioxidant defense components, with catalases in particular being more expressed in RPE/LULS53. Genes associated with iron homeostasis, such as *Ferritin HCH* and *IRP*, also displayed higher expression in RPE/LULS53, whereas *HO* showed slightly higher expression in RPE/LULS57 (20.29 vs. 15.50 TPM).

**Fig 4.**
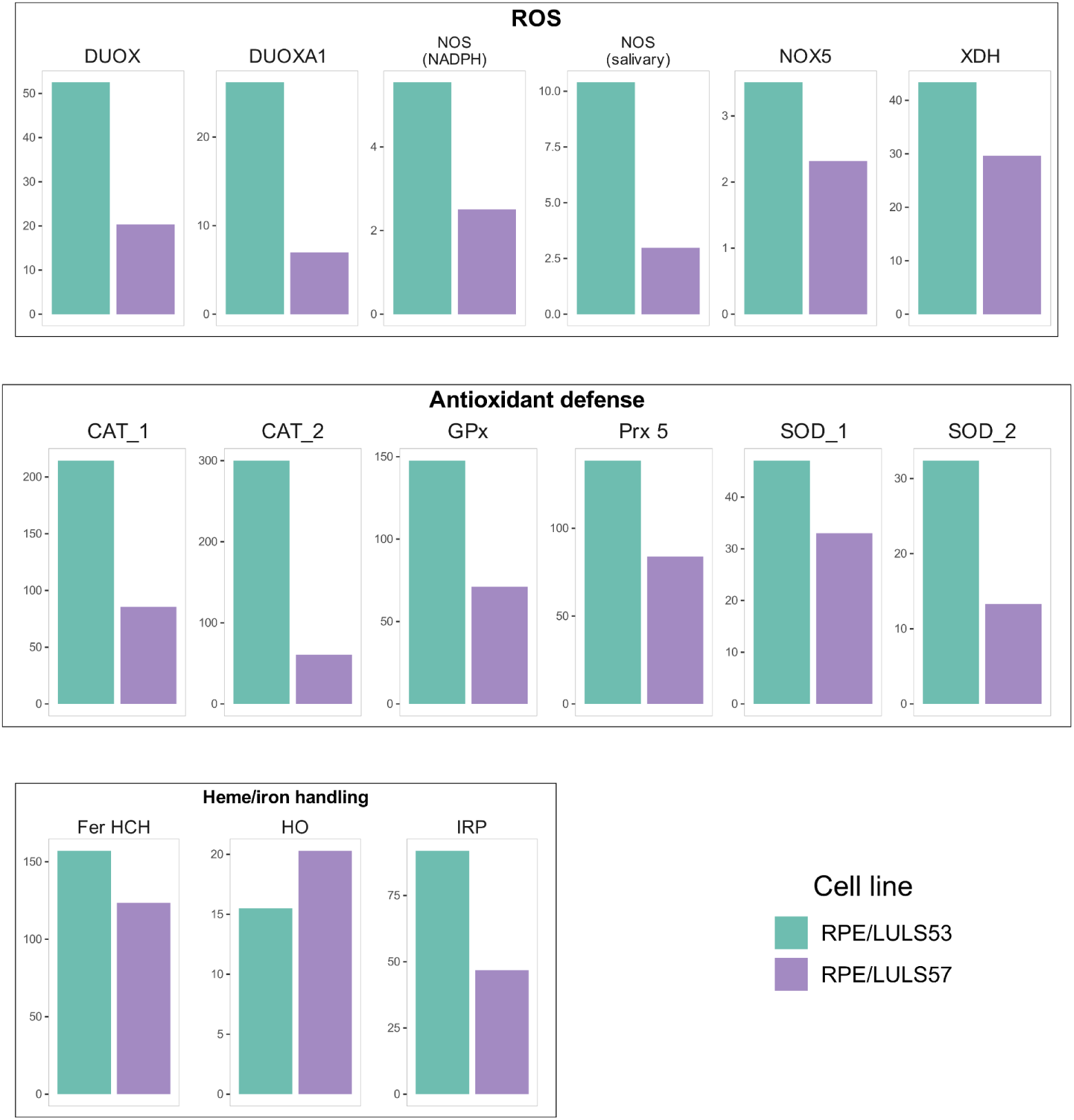
Expression of genes associated with the integrated redox homeostasis mechanism in RPE/LULS53 and RPE/LULS57 cell lines. Expression levels of genes involved in the integrated redox homeostasis mechanism are shown. Genes are grouped according to their roles in ROS metabolism, antioxidant defense, and heme/iron handling. Gene expression values were derived from RNA-seq data and normalized by TPM. Bars represent transcript abundance displayed for each gene on an independent TPM scale (y-axis).

Together, these patterns suggest that the RPE/LULS53 cell line may possess a more active or regulated redox mechanism compared with RPE/LULS57. The complete dataset of this analysis with the expression levels is available in S7 Table. Additional components of these pathways, including gene fragments or more specific variants, were also examined and are presented in S3 Fig.

### *Rhodnius* embryo-derived cell lines express both resident and horizontally transmitted transposable elements

Transposable elements (TEs) and other repetitive sequences make up a large fraction of animal and plant genomes and their deregulation has been associated with a range of genetic diseases in humans^33^. In *Rhodnius*, TEs are estimated to represent approximately 19-23% of the genome, with elements of the *Mariner* family being the most abundant class^13,34–36^. While the mechanism preventing TE mobilization has been dissected in great detail in the fruit fly *D. melanogaster*, much less is known in more basal insect species such as triatomine insects. Therefore, the RPE/LULS53 and RPE/LULS57 cell lines will likely help to shed light on these mechanisms.

Hence, we investigated transposon expression by detecting transcripts derived from multiple TE classes in both cell lines (Fig. 5). Notably, sequences classified as Unknown represent the most abundant category of TE-related transcripts in both RPE/LULS53 and RPE/LULS57, reaching approximately 302,726 and 400,355 TPM, respectively. Among annotated elements, DNA transposons of the TcMar (*Mariner*) superfamily were the most highly expressed, with transcript levels of ≈50,394 TPM in RPE/LULS53 and ≈90,658 TPM in RPE/LULS57, consistent with previous reports in this species^13,34–36^. Other DNA transposons, such as the rolling-circle *Helitron* and *hAT* superfamilies were also highly expressed, with values ranging from ≈14,559 to 34,287 TPM across the two cell lines. Retrotransposons were also strongly represented. Several LINE elements, including *I*, *R1*, and *RTE*, displayed substantial expression levels in both cell lines. Among LTR retrotransposons, *Gypsy* constituted the most abundant family, whereas other elements such as *Pao* and *Copia* were detected at lower levels.

**Fig 5.**
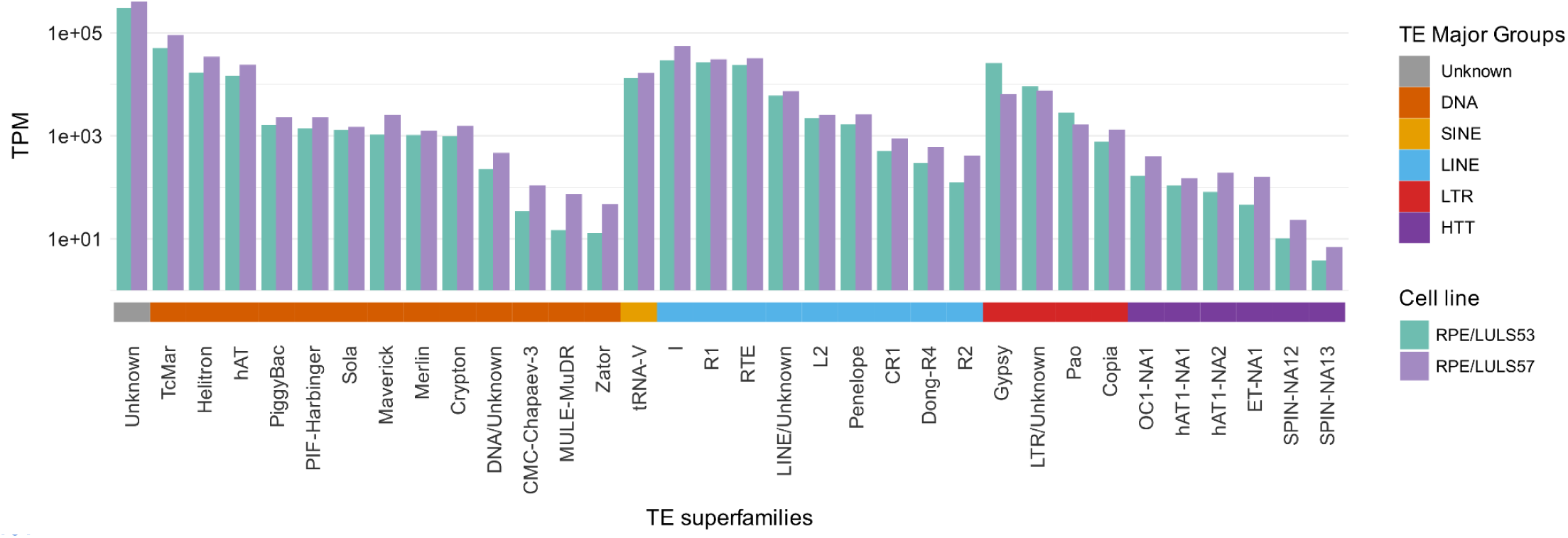
Expression profiles of transposable elements in the RPE/LULS53 and RPE/LULS57 cell lines. Bar plots showing TE expression levels detected in RPE/LULS53 and RPE/LULS57 cell lines. TE families are displayed along the x-axis and grouped by classification, as indicated by the colored bar below. Expression levels are normalized by TPM and displayed on a base-10 logarithmic scale to accommodate data covering multiple orders of magnitude.

Although most TE families were expressed in both cell lines, quantitative differences were observed between them. For many DNA transposons - including *TcMar* and *Helitron* - and LINE elements - such as *I* and *RTE* - transcript levels were consistently higher in RPE/LULS57 compared with RPE/LULS53. In contrast, certain elements such as *Gypsy* displayed markedly higher expression in RPE/LULS53 (≈25,816 TPM) than in RPE/LULS57 (≈6,437 TPM).

Interestingly, in addition to resident transposons - which have resided in the *Rhodnius* genome for longer evolutionary periods and are vertically transmitted across generations - it was shown that this insect species acquired new transposable elements by horizontal transfer over the past five million years. These horizontally transmitted transposons (HTTs) include *OposCharlie1* (*OC1*), *hAT1*, *ExtraTerrestrial* (*ET*) and *SPACE INVADERS* (*SPIN*) elements originating from vertebrate hosts, such as opossums, squirrel monkeys and bats and likely invaded the *Rhodnius* genome during blood feeding^13,37^.

In this context, we quantified their expression levels in the *R. prolixus* cell lines. Transcripts derived from HTTs were detected, although at substantially lower levels (from ≈3-394 TPM across different elements in both cell lines) compared to highly expressed resident transposons, such as *Mariner* and *Gypsy*. HTT transcript levels were consistently higher in RPE/LULS57, with values 1.3 to 3.4 times greater than in RPE/LULS53. The complete dataset of TE expression levels is available in S8 Table. Collectively, these results indicate that both embryo-derived cell lines express a diverse repertoire of resident and horizontally transferred transposable elements.

### The *Rp-β-Tubulin* promoter drives eGFP expression in the RPE/LULS57 cell line

Finally, we evaluated whether our transcriptomic data could support the development of new tools for genetic studies in *Rhodnius* embryo-derived cell lines. To this end, we aimed to generate a recombinant expression vector capable of driving eGFP expression in cell lines. The identification of a suitable promoter requires not only high transcriptional activity, but also a well-resolved genomic region. Although the sequencing and assembly of the *Rhodnius* genome (RproC3) have greatly advanced genetic and genomic studies, a substantial number of genes still present incomplete annotation or poorly assembled upstream regions^38^.

Based on these criteria, the *Rp-β-Tubulin* (*Rp-β-Tub*) gene, encoding the β-tubulin chain, was selected. This gene ranks among the most highly expressed in the *R. prolixus* cell lines, with TPM values of 974.39 in RPE/LULS53 and 505.87 in RPE/LULS57 (S2 and S3 Tables). Although other genes displayed higher expression levels, such as *Rp-α-Tubulin* (TPM = 996.05 in RPE/LULS57), *Rp-β-Tub* was prioritized because its upstream genomic region encompassing the putative promoter seemed to be well sequenced and assembled.

In agreement with this, we were able to PCR-amplify a ≈500 bp fragment upstream of the *Rp-β-Tub* coding sequence spanning the putative promoter region, and clone it upstream of an eGFP coding sequence in a pBluescript-derived plasmid^39^. This approach yielded the final pBlue-*Rp-β-Tub*-eGFP reporter construct (Fig. 6a). Transient transfection of this construct into RPE/LULS57 cells resulted in a clearly detectable eGFP signal three days post-transfection, as per immunofluorescence assays using anti-GFP antibodies (Fig. 6b). As expected for a freely diffusible protein, eGFP fluorescence was observed both in the nucleus and in the cytoplasm of the cells.

**Fig 6.**
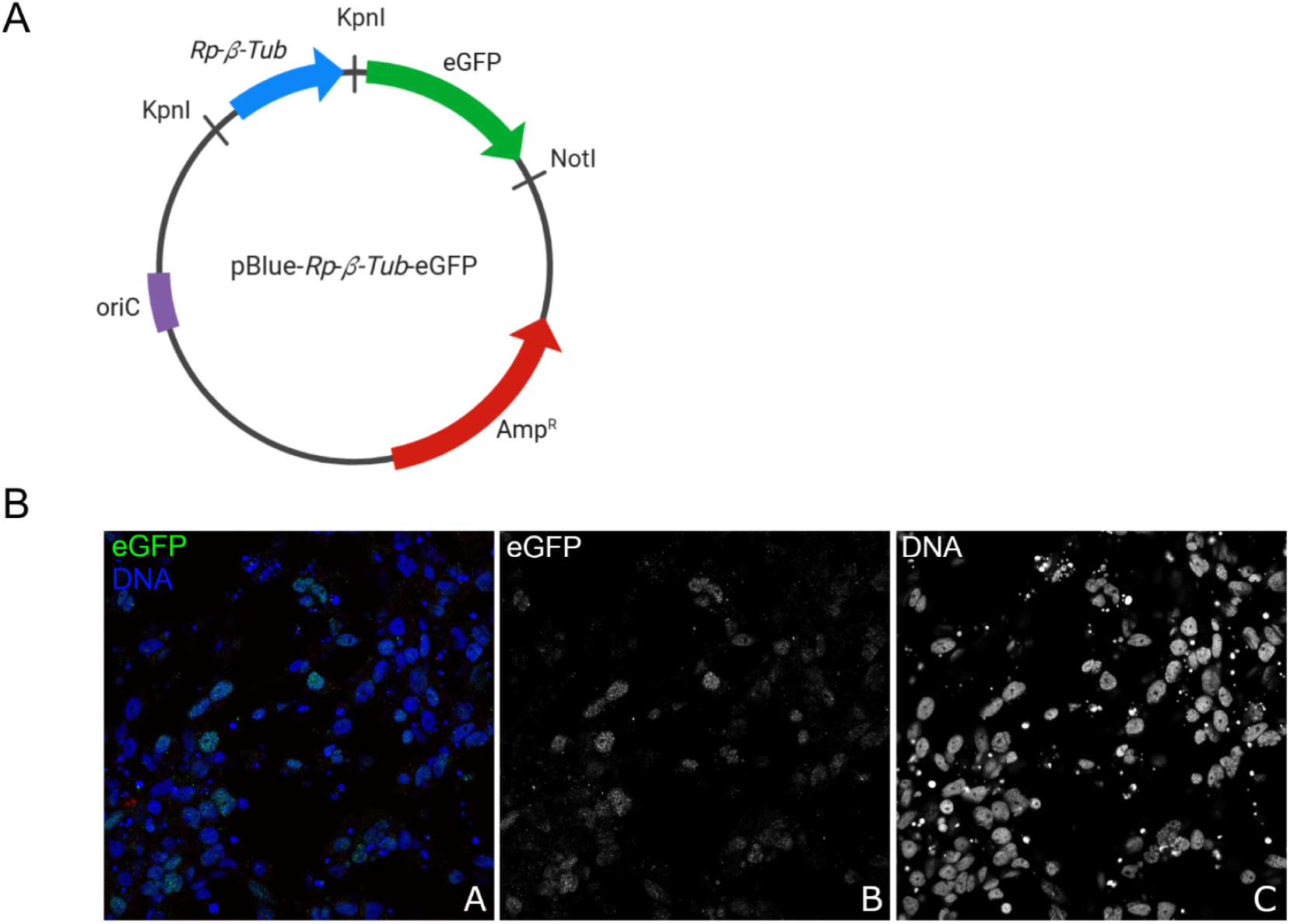
Expression of an eGFP reporter sequence driven by the *Rp-β-tub* promoter in RPE/LULS57 cells. (A) Schematic representation of the recombinant plasmid pBlue-*Rp-β-Tub*-eGFP. A ≈500 bp fragment corresponding to the *Rp-β-Tub* promoter was cloned upstream of an eGFP sequence encoding cDNA. The restriction sites KpnI and NotI used for cloning are indicated. Arrows indicate the direction of transcription for each element. (B) Immunofluorescence staining using a specific antibody against eGFP reveals a clear signal in both the cytoplasm and nuclei of RPE/LULS57 cells (A, green; B, single-channel). Nuclear DNA was counterstained with DAPI (A, blue; C, single-channel).

Thus, our transcriptomic datasets provide a reference framework for the design of expression systems, enabling the expression of chimeric proteins in *R. prolixus* embryo-derived cell lines.

## Discussion

RPE/LULS53 and RPE/LULS57 are stable cell lines derived from embryos of *R. prolixus*, a hematophagous hemipteran insect, and provide experimentally amenable systems to investigate conserved developmental signaling pathways, innate immune responses, non-coding RNA biology and genome stability in a medically relevant insect vector. The recent establishment of an additional *Rhodnius* cell line^40^ highlights the growing availability of *in vitro* systems for triatomine research and the emerging opportunities to investigate vector biology. Here, we provide the first comprehensive RNA-seq-based characterization of RPE/LULS53 and RPE/LULS57, representing an important step towards expanding the study of triatomine insects.

Our global transcriptomic analyses revealed that both cell lines share a substantial fraction of expressed genes, reflecting a conserved core transcriptional program consistent with their common embryonic origin. This includes highly expressed genes involved in processes such as translation, cytoskeletal organization and mitochondrial function, exemplified by elongation factor 1α, tubulins and cytochrome c oxidase subunits. However, the presence of cell line-specific transcripts and qualitative differences among the most abundant genes likely reflect the existence of two divergent phenotypes. This divergence may result from intrinsic variability amongst the multiple embryos from which each cell line was generated, as well as adaptation to distinct culture conditions and establishment timelines^21^. Therefore, each cell line may be preferentially suited for specific evolutionary and functional investigations, depending on the specific gene or biological process of interest, despite both having been derived from embryonic tissues using similar dissociation and culture procedures.

Consistent with this, we identified 391 cell line-specific genes in RPE/LULS53. Among them, one of the most highly expressed genes was *Cuticular protein 100a*, which encodes a structural component of the chitin-based cuticle and has been associated with cuticle development during embryonic and pupal stages in other insects^41^. On the other hand, the RPE/LULS57 cell line expressed 1,088 specific genes. Among them, we found *Neuroligin-2*, which encodes a synaptic cell-adhesion protein involved in the organization of inhibitory (GABAergic) synapses. The dysregulation of this protein in humans has been associated with neurodevelopmental and neuropsychiatric conditions, including autism spectrum disorders, schizophrenia, and anxiety-related phenotypes^42^.

This cell line also expressed precursors of specific miRNAs, such as miR-2c, miR-71 and miR-13b, which were present at much lower or undetectable levels in the RPE/LULS53 cell line. In total, we identified 29 miRNA precursors uniquely or preferentially expressed in RPE/LULS57 and, out of these, 12 have been previously reported in *R. prolixus*^17^. Among these, miR-71 has been implicated in the regulation of multiple processes across invertebrates, including development, stress responses and immune function^43^, whereas miR-275 has been associated with blood digestion and egg development in hematophagous insects^44^. Together, these findings indicate that the two cell lines also differ in the expression patterns of non-coding RNAs.

To further characterize the functional profiles of each cell line, we performed Gene Ontology enrichment and pathway-level analyses. While both cell lines retained core components, RPE/LULS53 was enriched for processes related to signal transduction, cellular response to stress, animal organ morphogenesis and protein serine/threonine kinase activity, whereas RPE/LULS57 was enriched for functions associated with DNA-directed RNA polymerase activity and nucleic acid conformation isomerase activity. These enrichment patterns indicate functional specialization within the two cell lines, suggesting that they may represent distinct cellular phenotypes rather than merely technical variants of the same embryonic origin.

In addition, how the insect deals with the reactive oxygen species released from the metabolism of heme is an area of active investigation and the *R. prolixus* cell lines might provide an important tool to further dissect this important process. Considering this, we observed that genes involved in redox homeostasis were predominantly upregulated in RPE/LULS53, suggesting a more active redox mechanism, potentially linked to differences in metabolic activity or stress responsiveness. This result is consistent with the enrichment of Gene Ontology terms related to cellular stress responses in this cell line, reinforcing the idea that RPE/LULS53 may be better equipped to subsist to oxidative challenges. This may be particularly relevant in the context of hematophagy, where cells are exposed to high levels of heme-derived oxidative stress.

Thus, the choice of cell line should be guided by the specific biological question under investigation. To support such applications, we provide a comprehensive catalogue of genes, including their expression levels in each cell line and their annotated orthologs in other organisms (S2 and S3 Tables). This resource is expected to facilitate the design of future functional studies using *R. prolixus* cell lines.

Beyond protein-coding genes, the results also have implications for the study of small non-coding RNAs in *Rhodnius*. Our previous work has shown that *Rhodnius* ovaries and embryos express Piwi-interacting RNAs (piRNAs), as well as core components of the piRNA pathway^11–13^. These small RNAs associate with Piwi proteins and are involved in the downregulation of transposable and repetitive sequences in the genome of metazoans^45,46^. By silencing transposable elements, the pathway ensures the maintenance of genome stability. In agreement with this, both RPE/LULS53 and RPE/LULS57 express comparable levels of transcripts from a variety of transposons. Interestingly these include not only resident transposable elements that have populated the genome for long evolutionary timescales, but also horizontally transmitted transposons that have been acquired more recently from opossums, squirrel monkeys and bats^13,37^. Therefore, these cell lines will enable the dissection of molecular mechanisms that have been well characterized in *D. melanogaster* and a few other holometabolous insects, but remain largely unknown in hemimetabolous insect species such as *Rhodnius*.

Similarly, these systems will be pivotal to investigate the miRNA pathway, the antiviral response and the ternary interaction among triatomine insects, viruses and the Chagas disease agent *Trypanosoma cruzi*^12,14,17,47^. Although miRNAs in *R. prolixus* have begun to be characterized in recent studies, their regulatory roles remain only partially understood, highlighting the potential of these cell lines for further investigation.

Finally, functional studies will require the development of new genetic tools to enable the expression of specific factors, proteins or enzymes in *Rhodnius* cell lines. As a proof of concept, we demonstrate that the expression of genes or chimeric proteins can be achieved by harnessing information derived from our transcriptomic datasets. Specifically, we employed the promoter of the *Rp-β-tub* gene, which is highly expressed in both RPE/LULS53 and RPE/LULS57 cell lines, to drive the expression of an eGFP reporter cDNA in RPE/LULS57 cells. It is conceivable that the expression of a sequence of interest can be modulated through the selection of promoters derived from genes with varying expression levels. These systems could also be adapted to express chimeric proteins bearing eGFP or other molecular tags, enabling additional assays such as co-immunoprecipitation, chromatin immunoprecipitation and RNA immunoprecipitation, among others.

Therefore, by defining the gene expression landscapes of these two embryo-derived cell lines, this study establishes a foundational transcriptomic resource that complements *in vivo* analyses and facilitates future functional, comparative and evolutionary studies as well as mechanistic investigations of gene regulation in triatomines and other hemimetabolous insects.

## Author contributions

**Laura Tavares:** Data curation, Formal analysis, Investigation, Methodology, Validation, Visualization, Writing – original draft, Writing – review & editing. **Anna Garcia:** Supervision. **Lesley Bell-Sakyi**: Investigation. **Tarcísio Brito:** Data curation, Formal analysis, Methodology, Supervision, Visualization, Writing – original draft. **Attilio Pane:** Conceptualization, Data curation, Formal analysis, Funding acquisition, Investigation, Methodology, Project administration, Resources, Supervision, Validation, Visualization, Writing – original draft, Writing – review & editing.

## Data availability

Original datasets are available publicly at the Sequence Read Archive (SRA) database and are all part of the Bioproject XXXXXX.

## Acknowledgements

We are grateful to Bernardo Carvalho, Pedro Lagerblad de Oliveira and Marcia Cury El-Cheikh for their constant support. We also appreciate the support of members of the Functional Genomics Laboratory for helpful insights and critical reading of the manuscript. We are grateful to Dr. Flavio Alves Lara and Diego Augusto Souza Oliveira for technical assistance with *R. prolixus* cell lines. We thank the staff of the LaCTAD from State University of Campinas (UNICAMP) for library preparation and RNA-seq sequencing.

This work was supported by the National Council for Scientific and Technological Development (CNPq) (428100/2018-0; AP), the Carlos Chagas Filho Foundation for Research Support of the State of Rio de Janeiro (FAPERJ) (E-26/210.339/2024; AP) and the Brazilian Federal Agency for Support and Evaluation of Graduate Education (CAPES) (LT, AG and AP). The funders had no role in study design, data collection and analysis, decision to publish or preparation of the manuscript.

## Supporting information

**S1 Fig.**
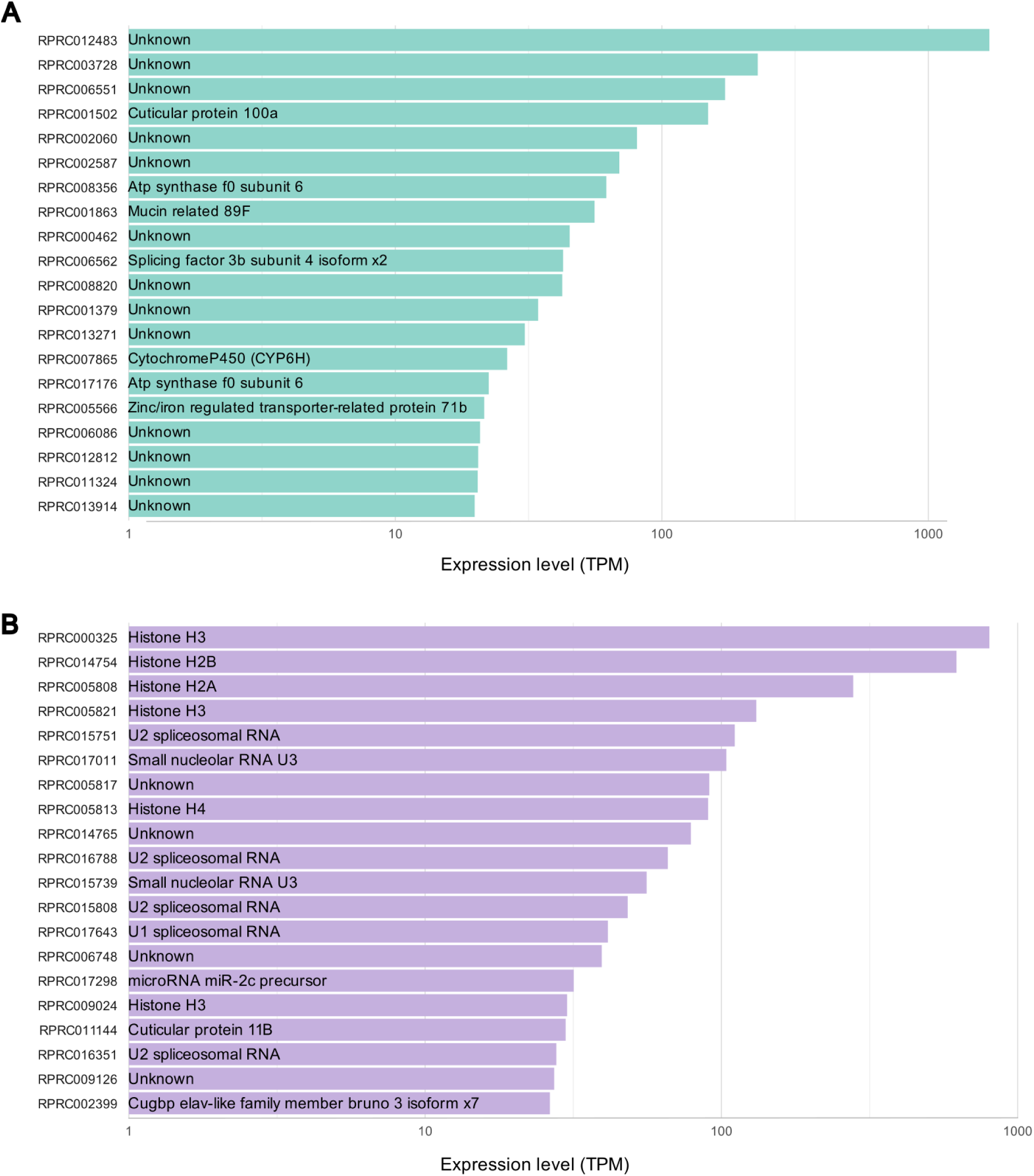
Top 20 transcripts most expressed exclusively in RPE/LULS53 and RPE/LULS57. (A-B) Bar plots showing the 20 most highly expressed transcripts among those exclusively expressed in RPE/LULS53 (A, green) and RPE/LULS57 (B, lilac), based on RNA-seq data. The x-axis represents the transcript abundance in TPM and the y-axis shows transcript IDs, with the corresponding functional annotation indicated within each bar.

**S2 Fig.**
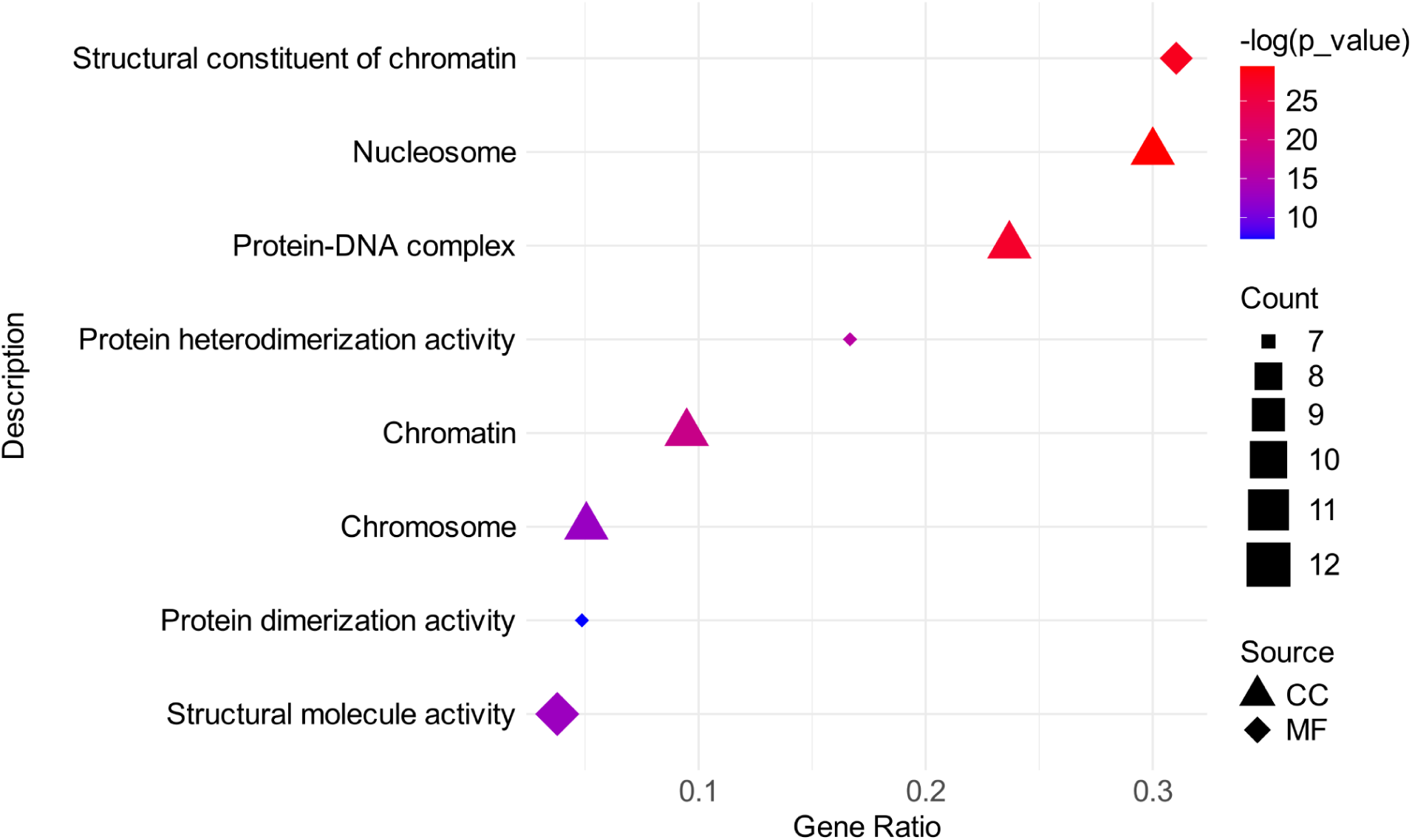
Gene Ontology enrichment analysis of genes exclusively expressed in RPE/LULS57. Dot plots showing significantly enriched GO terms for genes exclusively expressed in RPE/LULS57. Each point represents an enriched GO term, with the x-axis indicating the gene ratio (proportion of genes in the input list associated with the term) and the y-axis displaying the corresponding functional description. Point size reflects the number of genes associated with each term, while color indicates statistical significance in −log(p-value). Distinct shapes represent GO categories Cellular Component (CC, triangle) and Molecular Function (MF, diamonds).

**S3 Fig.**
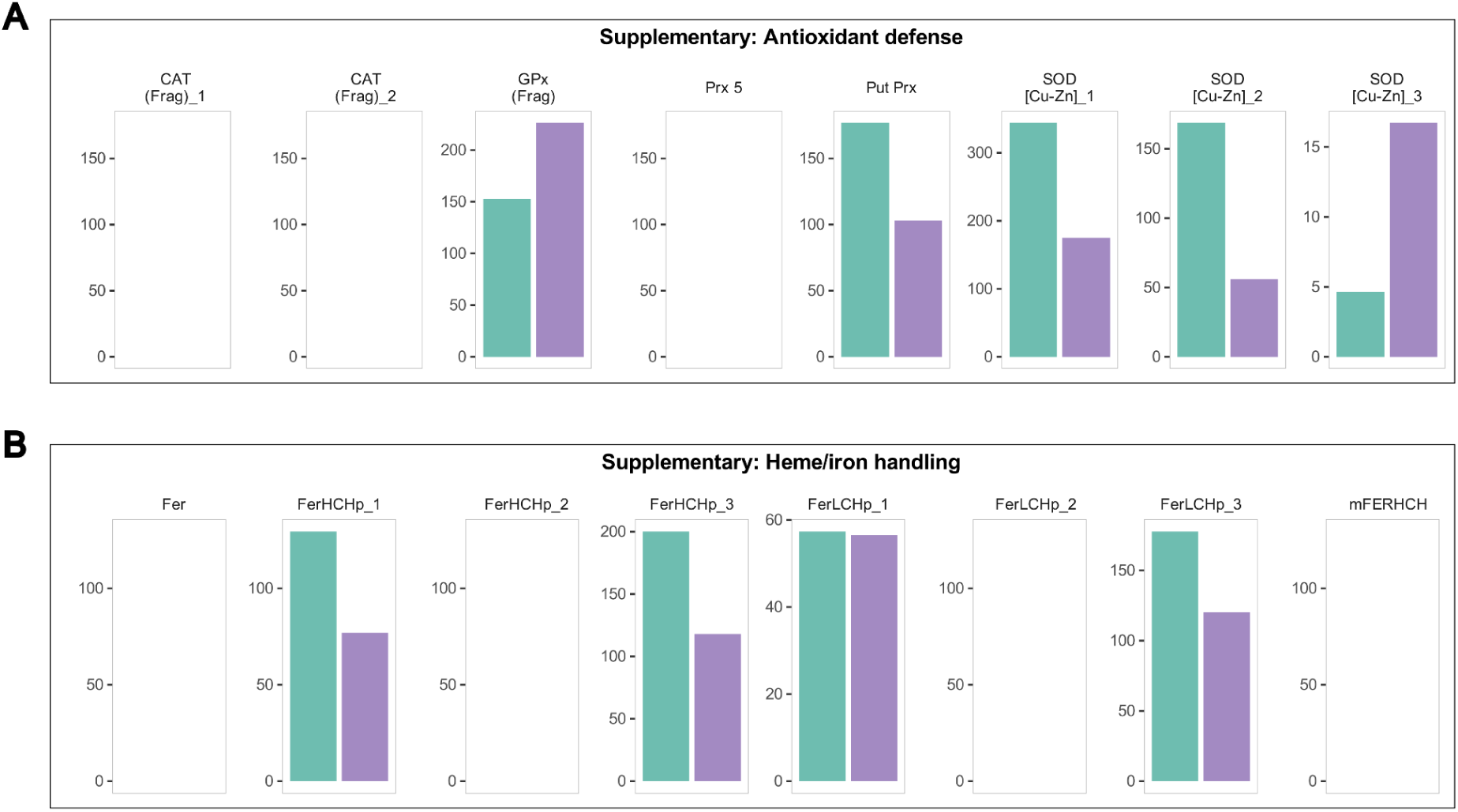
Expression of additional genes associated with the integrated redox homeostasis mechanism in RPE/LULS53 and RPE/LULS57 cell lines. Expression levels of additional genes involved in the integrated redox homeostasis mechanism are shown. Genes are grouped according to their roles in ROS metabolism and heme/iron handling. Gene expression values were derived from RNA-seq data and normalized by TPM. Bars represent transcript abundance displayed for each gene on an independent TPM scale (y-axis). RPE/LULS53 is shown in green and RPE/LULS57 in lilac.

**S1 Table. RNA-seq read processing and mapping statistics.** Total number of raw reads and after trimming and mapping against RproC3 genome RPE/LULS53 and RPE/LULS57 cell lines.

**S2 Table. Global gene expression in the RPE/LULS53 cell line.** All expressed genes with transcript abundance normalized as TPM, after applying the cutoff > 1. The table includes gene ID, TPM values and functional description. Annotations were derived from the official genome annotation, BLASTx results or g:Orth predictions - as described in the Methods section.

**S3 Table. Global gene expression in the RPE/LULS57 cell line.** All expressed genes with transcript abundance normalized as TPM, after applying the cutoff > 1. The table includes gene ID, TPM values and functional description. Annotations were derived from the official genome annotation, BLASTx results or g:Orth predictions - as described in the Methods section.

**S4 Table. Genes exclusively expressed in the RPE/LULS53 cell line.** All exclusively expressed genes in RPE/LULS53 normalized by TPM. The table includes gene ID, TPM values and functional description derived from the official genome annotation, BLASTx results or g:Orth predictions - as described in the Methods.

**S5 Table. Genes exclusively expressed in the RPE/LULS57 cell line.** All exclusively expressed genes in RPE/LULS57 normalized by TPM. The table includes gene ID, TPM values and functional description derived from the official genome annotation, BLASTx results or g:Orth predictions - as described in the Methods.

**S6 Table. TPM values of genes associated with developmental, stress, and immune signaling pathways in RPE/LULS53 and RPE/LULS57 cell lines.** Genes are grouped by the pathways ERK, JAK/STAT, Wnt, Notch, Hedgehog, Toll, Hippo, IMD, BMP and PCP.

**S7 Table. TPM values of genes associated with the integrated redox homeostasis mechanism in RPE/LULS53 and RPE/LULS57 cell lines.** Genes are grouped by the processes ROS metabolism, antioxidant defense, and heme/iron handling.

**S8 Table. TPM values of transposable element families in RPE/LULS53 and RPE/LULS57 cell lines.** Transposable elements are grouped into major classes (DNA, SINE, LINE, LTR, HTT and Unknown) and further classified by superfamily.

**S9 Table. Genes involved in major developmental and immune signaling pathways.** List of genes associated with ERK, JAK/STAT, Wnt, Notch, Hedgehog, Toll, Hippo, IMD, BMP and PCP signaling pathways in *Rhodnius prolixus*. Genes were compiled from the literature and used to explore the expression of these pathways in the RPE/LULS53 and RPE/LULS57 cell lines. Superscript numbers indicate the source from which specific genes were curated. “1” indicates genes that were described by Lavore and collaborators (Lavore et al., 2015). “2” indicates genes that were described by Salcedo-Porras and coworkers (Salcedo-Porras et al., 2019).

**S10 Table. Genes involved in redox homeostasis mechanism.** List of genes associated with ROS metabolism, antioxidant defense and heme/iron handling processes in *Rhodnius prolixus*. Genes were compiled from the literature and used to explore the expression of these pathways in the RPE/LULS53 and RPE/LULS57 cell lines. Superscript numbers indicate the source from which specific genes were curated. “1” indicates genes that were described by Gandara and collaborators (Ganadara et al., 2021). “2” indicates genes that were identified by DIAMOND Blast X.

## References

1. Pérez-Molina, J. A. & Molina, I. Chagas disease. Lancet 391, 82–94 (2018).

2. World Health Organization. Chagas disease (American trypanosomiasis). https://www.who.int/news-room/fact-sheets/detail/chagas-disease-(american-trypanosomiasis) (2026).

3. World Health Organization. Ending the neglect to attain the Sustainable Development Goals: a road map for neglected tropical diseases 2021–2030. (WHO, 2021).

4. Nunes-da-Fonseca, R., Berni, M., Tobias-Santos, V., Pane, A. & Araujo, H. M. Rhodnius prolixus: From classical physiology to modern developmental biology. Genesis 55, (2017).

5. Lange, A. B., Leyria, J. & Orchard, I. The hormonal and neural control of egg production in the historically important model insect, Rhodnius prolixus: A review, with new insights in this post-genomic era. Gen. Comp. Endocrinol. 321–322, 114030 (2022).

6. Azambuja, P. et al. Rhodnius prolixus: from physiology by Wigglesworth to recent studies of immune system modulation by Trypanosoma cruzi and Trypanosoma rangeli. J Insect Physiol 97, 45–65 (2017).

7. Wigglesworth, V. B. The Physiology of Ecdysis in *Rhodnius prolixus* (Hemiptera). II. Factors controlling Moulting and ‘Metamorphosis’. J Cell Sci s2-77, 191–222 (1934).

8. Beament, J. W. L. The Formation and Structure of the Chorion of the Egg in an Hemipteran, Rhodnius prolixus. J Cell Sci s2-87, 393–439 (1946).

9. Kelly, G. M. & Huebner, E. Embryonic development of the hemipteran insect Rhodnius prolixus. Journal of Morphology 199, 175–196 (1989).

10. Lima, L. et al. Gene Editing in the Chagas Disease Vector by Cas9-Mediated ReMOT Control. CRISPR J 7, 88–99 (2024).

11. Brito, T., et al. Transcriptomic and functional analyses of the piRNA pathway in the Chagas disease vector Rhodnius prolixus. PLoS Negl. Trop. Dis. 12, e0006760 (2018).

12. Brito, T. F. de et al. Transovarial transmission of a core virome in the Chagas disease vector Rhodnius prolixus. PLoS Pathog. 17, e1009780 (2021).

13. de Brito, T. F. et al. Embryonic piRNAs target horizontally transferred vertebrate transposons in assassin bugs. Front Cell Dev Biol 12, 1481881 (2024).

14. Cardoso, M. A. et al. The Neglected Virome of Triatomine Insects. Frontiers in Tropical Diseases vol. 3 Preprint at 10.3389/fitd.2022.828712 (2022).

15. Benrabaa, S. A. M., Orchard, I. & Lange, A. B. The role of ecdysteroid in the regulation of ovarian growth and oocyte maturation in Rhodnius prolixus, a vector of Chagas disease. J Exp Biol 225, (2022).

16. Medeiros, M. N. et al. Transcriptome and gene expression profile of ovarian follicle tissue of the triatomine bug Rhodnius prolixus. Insect Biochem. Mol. Biol. 41, 823–831 (2011).

17. Santiago, P. B. et al. Insights into the microRNA landscape of Rhodnius prolixus, a vector of Chagas disease. Sci Rep 13, 13120 (2023).

18. Mesquita, R. D. et al. Genome of Rhodnius prolixus, an insect vector of Chagas disease, reveals unique adaptations to hematophagy and parasite infection. Proc. Natl. Acad. Sci. U. S. A. 112, 14936–14941 (2015).

19. Cardoso, J. C. et al. Analysis of the testicle’s transcriptome of the Chagas disease vector Rhodnius prolixus. bioRxiv (2019) doi:10.1101/616193.

20. Marchant, A. et al. Comparing de novo and reference-based transcriptome assembly strategies by applying them to the blood-sucking bug Rhodnius prolixus. Insect Biochem. Mol. Biol. 69, 25–33 (2016).

21. Penrice-Randal, R. et al. New Cell Lines Derived from Laboratory Colony Do Not Harbour Triatoma Virus. Insects 13, (2022).

22. Schneider, I. Cell lines derived from late embryonic stages of Drosophila melanogaster. J Embryol Exp Morphol 27, 353–365 (1972).

23. Strunov, A. et al. Ultrastructural analysis of mitotic Drosophila S2 cells identifies distinctive microtubule and intracellular membrane behaviors. BMC Biol 16, 68 (2018).

24. Pan, M.-H., et al. Establishment and characterization of an ovarian cell line of the silkworm, Bombyx mori. Tissue and Cell 42, 42–46 (2010).

25. Marti, G. A. et al. Can Triatoma virus inhibit infection of Trypanosoma cruzi (Chagas, 1909) in Triatoma infestans (Klug)? A cross infection and co-infection study. J. Invertebr. Pathol. 150, 101–105 (2017).

26. Maria Vasconcelos Queiroz, A., et al. Antibodies response induced by recombinant virus-like particles from Triatoma virus and chimeric antigens from Trypanosoma cruzi. Vaccine 39, 4723–4732 (2021).

27. An, W., Yan, Y. & Ye, K. High resolution landscape of ribosomal RNA processing and surveillance. Nucleic Acids Res 52, 10630–10644 (2024).

28. Gasic, I. Regulation of Tubulin Gene Expression: From Isotype Identity to Functional Specialization. Front. Cell Dev. Biol. 10, 898076 (2022).

29. Akopian, D., Shen, K., Zhang, X. & Shan, S.-O. Signal recognition particle: an essential protein-targeting machine. Annu Rev Biochem 82, 693–721 (2013).

30. Temporal Regulation of Signal Recognition Particle During Translation. Journal of Molecular Biology 437, 169482 (2025).

31. Mabin, J. W., Lewis, P. W., Brow, D. A. & Dvinge, H. Human spliceosomal snRNA sequence variants generate variant spliceosomes. RNA 27, 1186–1203 (2021).

32. Graça-Souza, A. V., et al. Adaptations against heme toxicity in blood-feeding arthropods. Insect Biochemistry and Molecular Biology 36, 322–335 (2006).

33. Dong, W. et al. Transposons disrupt genomic stability and trigger cancers. Front. Cell. Infect. Microbiol. 15, 1704405 (2025).

34. Filée, J., Rouault, J.-D., Harry, M. & Hua-Van, A. Mariner transposons are sailing in the genome of the blood-sucking bug Rhodnius prolixus. BMC Genomics 16, 1061 (2015).

35. Fernández-Medina, R. D., Granzotto, A., Ribeiro, J. M. & Carareto, C. M. A. Transposition burst of mariner-like elements in the sequenced genome of Rhodnius prolixus. Insect Biochem Mol Biol 69, 14–24 (2016).

36. Montiel, E. E., Panzera, F., Palomeque, T., Lorite, P. & Pita, S. Satellitome Analysis of Rhodnius prolixus, One of the Main Chagas Disease Vector Species. Int. J. Mol. Sci. 22, 6052 (2021).

37. Gilbert, C., Schaack, S., Pace, J. K., 2nd, Brindley, P. J. & Feschotte, C. A role for host-parasite interactions in the horizontal transfer of transposons across phyla. Nature 464, 1347–1350 (2010).

38. Coelho, V. L. et al. Analysis of ovarian transcriptomes reveals thousands of novel genes in the insect vector Rhodnius prolixus. Sci. Rep. 11, 1918 (2021).

39. Pane, A., Jiang, P., Zhao, D. Y., Singh, M. & Schüpbach, T. The Cutoff protein regulates piRNA cluster expression and piRNA production in the Drosophila germline. EMBO J. 30, 4601–4615 (2011).

40. Hartley, C., Khoo, J. J., Darby, A., Makepeace, B. L. & Bell-Sakyi, L. New tick and insect cell line resources for vector-borne disease research from the Tick Cell Biobank. Rev. Rev. Elev. Med. Vet. Pays Trop. 78, 1–10 (2025).

41. Chen J, et al. Roles of a newly lethal cuticular structural protein, AaCPR100A, and its upstream interaction protein, G12-like, in Aedes aegypti. International Journal of Biological Macromolecules 268, 131704 (2024).

42. Zeppillo, T. et al. Functional Neuroligin-2-MDGA1 interactions differentially regulate synaptic GABAARs and cytosolic gephyrin aggregation. Communications Biology 7, 1157 (2024).

43. Naidoo, D., Brennan, R. & de Lencastre, A. Conservation and Targets of miR-71: A Systematic Review and Meta-Analysis. Noncoding RNA 9, (2023).

44. Bryant, B., Macdonald, W. & Raikhel, A. S. microRNA miR-275 is indispensable for blood digestion and egg development in the mosquito Aedes aegypti. Proceedings of the National Academy of Sciences 107, 22391–22398 (2010).

45. Santos, D. et al. What Are the Functional Roles of Piwi Proteins and piRNAs in Insects? Insects 14, (2023).

46. Onishi, R., Yamanaka, S. & Siomi, M. C. piRNA- and siRNA-mediated transcriptional repression in Drosophila, mice, and yeast: new insights and biodiversity. EMBO Rep 22, e53062 (2021).

47. Paes, M. C. et al. Gene expression profiling of Trypanosoma cruzi in the presence of heme points to glycosomal metabolic adaptation of epimastigotes inside the vector. PLoS Negl Trop Dis 14, e0007945 (2020).

48. Bhardwaj, V. et al. snakePipes: facilitating flexible, scalable and integrative epigenomic analysis. Bioinformatics 35, 4757–4759 (2019).

49. Martin, M. Cutadapt removes adapter sequences from high-throughput sequencing reads. EMBnet.journal 17, 10–12 (2011).

50. Dobin, A. et al. STAR: ultrafast universal RNA-seq aligner. Bioinformatics 29, 15–21 (2012).

51. Jin, Y., Tam, O. H., Paniagua, E. & Hammell, M. TEtranscripts: a package for including transposable elements in differential expression analysis of RNA-seq datasets. Bioinformatics 31, 3593–3599 (2015).

52. Buchfink, B., Reuter, K. & Drost, H.-G. Sensitive protein alignments at tree-of-life scale using DIAMOND. Nature Methods 18, 366–368 (2021).

53. Kolberg, L. et al. g:Profiler—interoperable web service for functional enrichment analysis and gene identifier mapping (2023 update). Nucleic Acids Res 51, W207–W212 (2023).

## References

1. Lavore A, et al. Comparative analysis of zygotic developmental genes in *Rhodnius prolixus* genome shows conserved features on the tracheal developmental pathway. Insect Biochemistry and Molecular Biology 64, 32–43 (2015).

2. Salcedo-Porras, N., Guarneri, A., Oliveira, P. L. & Lowenberger, C. *Rhodnius prolixus*: Identification of missing components of the IMD immune signaling pathway and functional characterization of its role in eliminating bacteria. PLOS ONE 14, e0214794 (2019).

3. Gandara, A. C. P. et al. ‘Urate and NOX5 Control Blood Digestion in the Hematophagous Insect *Rhodnius prolixus*’. Front. Physiol. 12, 633093 (2021).

